# Effect of p-benzoquinone, a cigarette smoke-derived component, on the architecture and function of human RBC

**DOI:** 10.1101/2024.10.21.619387

**Authors:** Neha Yadav, Santosh Kumar Mondal, Amit Kumar Mandal

## Abstract

p-benzoquinone (pBQ), a cigarette smoke-derived component, stands as one of the lethal oxidants of cigarette smoke. Health hazards caused by cigarette smoking are a profound global health concern. It has been reported to contribute significantly to several health disorders, such as lung cancer, chronic obstructive pulmonary disease (COPD), cardiovascular diseases, etc. These adverse health effects might be attributed to the complex mixture of approximately 4,000 chemicals present in cigarette smoke, such as nicotine, tar, nitric oxide, and p-benzosemiquinone (pBSQ). pBSQ, ranging from 100-200 µg per cigarette, gets oxidized to pBQ in the lungs of smokers and subsequently enters into the bloodstream. In the present study, we investigated the detrimental effect of pBQ on the architecture of human RBCs. Previously, we showed that the covalent modification of human hemoglobin with pBQ results in an alteration in its structure and function. In addition, we explored the potential neutralizing role of N-acetyl cysteine (NAC) in the reactivity of pBQ. Our results demonstrated that pBQ exposure to human red blood cells (RBCs), *ex vivo*, significantly decreases the reduced glutathione (GSH) content of RBCs, leading to the elevation of reactive oxygen species (ROS) and consequently causing oxidative damage to cellular components. pBQ exposure resulted in increased RBC sedimentation, altered membrane fluidity, enhanced lipid peroxidation, and significant morphological changes, such as bulging and distortion on the surface of RBCs, which eventually compromise membrane integrity. In addition, the coexistence of NAC with pBQ was found to be effective in neutralizing pBQ, thereby restoring GSH levels and RBC membrane architecture. These findings highlight the potential of NAC in mitigating the harmful effects of cigarette smoke on RBCs.

**Graphical abstract:** 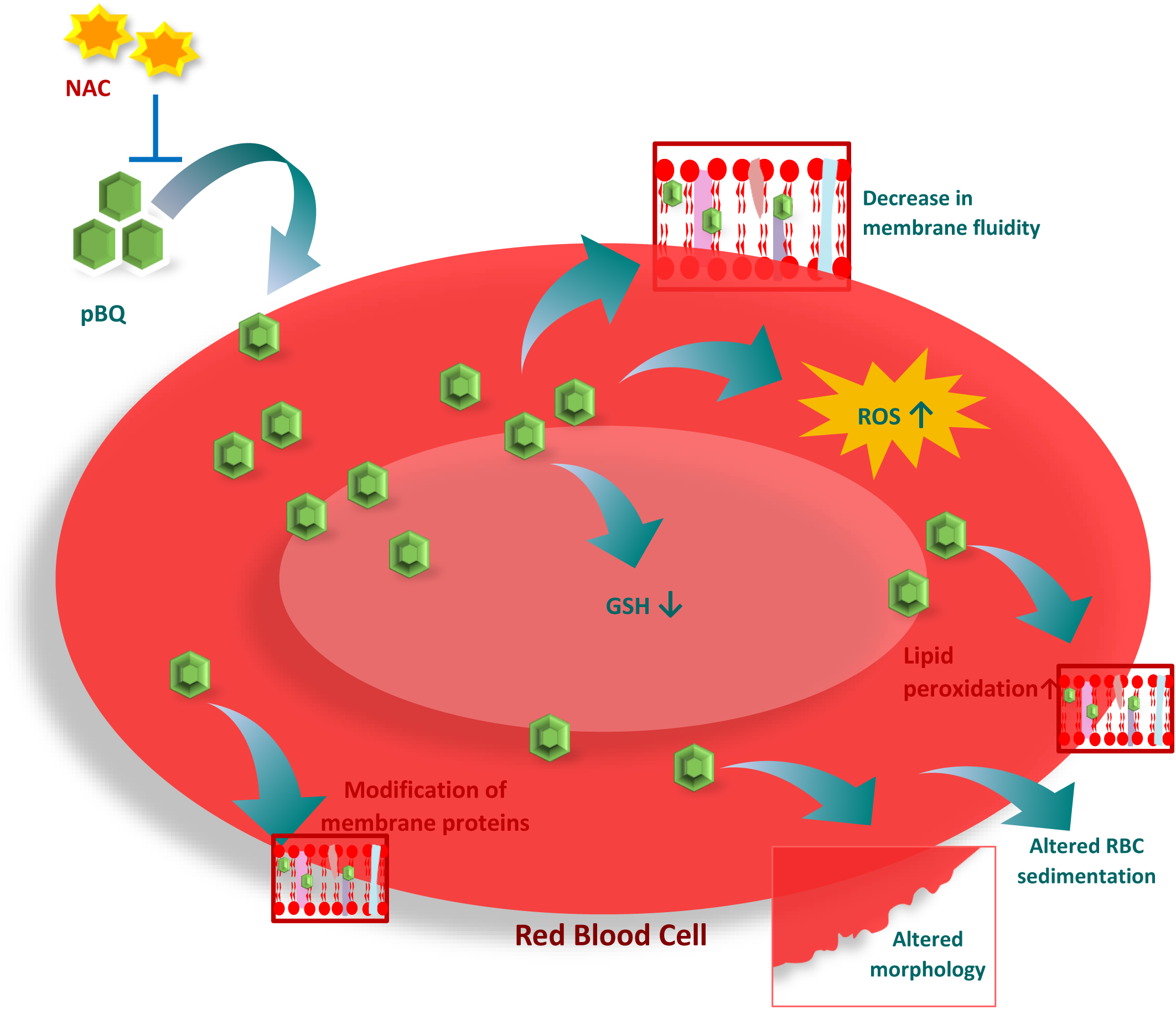

## Introduction

Cigarette smoking is the primary cause to blame for a wide range of illnesses and is one of the major causes of death globally. Cigarette smoking causes several life-threatening diseases, such as lung cancer, chronic obstructive pulmonary disease (COPD), stroke, cardiovascular diseases, etc., which eventually cause premature death [1]. Most of these tobacco-smoking fatalities occur in underdeveloped countries, and statistical data reveal that smokers pass away 14 years sooner than non-smokers. Tobacco is responsible for around 3 million deaths worldwide yearly; by 2030, this number is expected to rise to 10 million. In India, 14% of women and 47% of men smoke cigarettes regularly [2]. Cigarette smoke (CS) is a complex mixture of approximately 4000 chemicals such as nicotine, tar, carbon monoxide, nitric oxides, heavy metals (cadmium, chromium, selenium, etc), polycyclic aromatic hydrocarbons, and several oxidants like nitric oxide, carbon monoxide, p-benzosemiquinone. In commercially available cigarettes, p-benzosemiquinone content is in the range of 100–200 µg [3–6]. It has been reported that quinones have toxic and hazardous effects, such as oxidative damage and the formation of adducts with cellular components like proteins, DNA, etc. In the lungs of smokers, p-benzosemiquinone of CS gets oxidized to p-benzoquinone (pBQ) and enters into the bloodstream. pBQ can cross the red blood cell (RBC) membrane and lead to irreversible posttranslational modification of human hemoglobin through covalent bonding with Cys93 of β hemoglobin that ultimately alters its structure and function [7,8].

pBQ has already been reported to be linked with Cigarette smoking-induced health disorders, including myelodysplastic syndromes (MDS), myocardial injury, and cardiomyocyte damage [9,10]. The exposure of RBCs from CS-derived oxidants causes alterations in the stability of the membrane [11]. Gangopadhyay *et al.* reported a decrease in the RBC membrane fluidity in COPD patients, which might be associated with cigarette smoking [12]. However, the screening of the cigarette smoke components as a causative factor for the above changes has not been explored yet.

pBQ, a strong oxidant derived from CS, can modify cellular components like proteins through covalent bonding with free and accessible cysteine residues. It can also generate reactive oxygen species (ROS), which might lead to oxidative stress and, subsequently, oxidative modifications of cellular components [8,13,14]. The rise in the level of ROS can be accompanied by a decrease in the level of antioxidants. In the present study, we investigated the effect of pBQ on different components of RBCs, *ex vivo*, such as reduced glutathione (GSH), ROS production, membrane fluidity, and morphology of RBC. Additionally, we studied the efficacy of N-acetyl cysteine (NAC) in neutralizing the effect of pBQ.

## Materials and methods

### Materials

p-benzoquinone, N-acetyl cysteine, 1,6-Diphenyl-1,3,5-hexatriene, 2′,7′-Dichlorofluorescein diacetate, 2-Thiobarbituric acid, and 5,5′-dithiobis-(2-nitrobenzoic acid), Dimethyl sulfoxide (DMSO) was purchased from Sigma Aldrich. LC/MS grade water and formic acid were obtained from Thermo Fisher. All used chemicals were of analytical grade.

### Preparation of RBC suspension

2 ml venous blood sample was collected in vacutainer tubes coated with ethylenediamine tetraacetic acid (EDTA) from AIIMS, Kalyani. Plasma was removed by centrifugation of the blood sample at 3000 rpm for 10 min at 25 °C. The resulting packed RBCs were washed thrice with Phosphate Buffer Saline (PBS), pH 7.4. RBCs (10% hematocrit) were incubated with pBQ at various concentrations (0.25 mM, 0.5 mM, 0.75 mM, 1 mM) in different sets for 4 h at 37 °C.

### GSH estimation

GSH levels within RBCs were quantified following a modified protocol outlined by Mandal *et al.* [15]. Briefly, RBCs (10% hematocrit) were exposed to varying concentrations of pBQ for 4 h at 37 °C as mentioned above and washed thrice with PBS, pH 7.4, by centrifugation at 3000 rpm for 10 min. Proteins were precipitated by adding four-fold 5% meta-phosphoric acid to the RBCs, followed by vortexing and centrifugation at 6000 rpm for 10 min at 4 °C. The resulting supernatant was subjected to treatment with 100 µM 5,5′-dithiobis-(2-nitrobenzoic acid) (DTNB), dissolved in 0.1 M KP (Potassium phosphate) buffer containing 10 mM EDTA, pH 7.4, for 10 minutes, and the newly generated 2-nitrobenzoic acid (TNB) was quantified at 412 nm using a UV-Vis spectrophotometer (Shimadzu, UV-1900i). The measured concentration of TNB is proportional to the concentration of GSH. To evaluate the impact of NAC in neutralizing the pBQ effect over GSH, RBCs were simultaneously exposed to 0.5 mM pBQ with 0.5 mM and 1 mM NAC in separate sets.

### ROS level measurement

ROS levels were measured by a little modification of the method described by Martínez-Vieyra *et al.* [16]. The aforementioned RBCs suspension incubated with varying concentrations of pBQ were washed thrice with PBS buffer, pH 7.4, by centrifuging at 3000 rpm for 10 min. Subsequently, RBCs were treated with 20 µM 2′,7′-Dichlorofluorescein diacetate (DCFDA) dissolved in DMSO for 1 h at 37 °C prior to the analysis using a fluorescence spectrophotometer (Fluoromax-3, Horiba Jobin Yvon). Upon entering into the RBCs, the diacetate group of DCFDA is cleaved by esterases present inside RBCs, and the resulting dichlorofluorescein (DCF) gives fluorescence by reacting with the free radicals generated within the RBCs. The fluorophore was excited at a wavelength of 485 nm, and emissions were recorded at 538 nm. The band width for both excitation and emission was maintained at 10 nm. To assess the efficacy of NAC in modulating ROS levels, RBCs were concurrently exposed to 0.5 mM pBQ with 0.5 mM and 1 mM NAC in separate sets. RBCs were subsequently incubated with DCFDA for 1 h at 37 °C and analysed using a fluorescence spectrophotometer.

The fluorescence data was corrected using the inner filter effect to alleviate the effect caused by the absorption of the light at the excitation wavelength and emission wavelength by the hemoglobin present within RBCs. The adjusted fluorescence was calculated using the following equation:

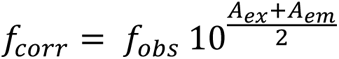

Where *f_corr_* and *f_obs_* represent the corrected and measured fluorescence intensities, respectively, while *A_ex_* and *A_em_* denote the absorbance measured at the excitation and emission wavelengths.

### RBC membrane fluidity

Membrane fluidity was measured using the Thompson method [17] with minor modifications. In brief, RBCs were exposed to ice-cold 5 mM KP buffer, pH 7.4, a hypotonic solution, followed by centrifugation at 10000 rpm for 10 mins at 4 ⁰C. The resulting pellet of erythrocytes was washed repeatedly until it became white in color, indicating the removal of the hemoglobin content of RBCs. Subsequently, the pellet containing RBCs devoid of hemoglobin (ghost RBCs) was subjected to wash with PBS buffer, pH 7.4, thrice and stored at 4°C until its use for further use in the experiment. The membrane protein content of the ghost RBCs was determined using the Bradford assay. 2 mg/ml of RBC membrane proteins were incubated with varying concentrations of pBQ. Following incubation, the membrane proteins were washed thrice with PBS buffer, pH 7.4, prior to fluorescence measurements. To assess the neutralizing impact of NAC on membrane fluidity, ghost RBCs were incubated at 0.5 mM pBQ in the presence of 0.5 mM and 1 mM NAC in separate sets.

1 mL of aforementioned ghost RBCs suspension containing 60 µg of RBC membrane proteins was incubated with 1,6-Diphenyl-1,3,5-hexatriene (DPH) to a final concentration of 3 µM for 1 h. The fluorescence measurements were taken in a fluorescence spectrophotometer (Hitachi F-7000) using 360 nm and 430 nm of excitation and emission wavelengths, respectively. The band width for excitation and emission was set to 10 nm, and the temperature of the system was maintained at 37 °C during the measurements. Fluorescence Anisotropy (r) was calculated using the following equation: r = (I_v_ − I_h_ × G)/(I_v_ − 2I_h_ × G), where I_v_ and I_h_ represent intensities measured through the polarizers oriented parallel and perpendicularly, respectively, to the vertical axis of the polarized excitation light, and G (I_v_/I_h_) is the correlation factor [18,19].

### RBC morphology

The impact of pBQ on the RBC morphology was investigated using a Field Emission Scanning Electron Microscope (FESEM) by following the protocol described by Etheresia Pretorius [20]. In brief, RBCs were exposed to 1 mM pBQ for 4 h, 37 °C, and were washed with PBS buffer, pH 7.4, by centrifuging at 3000 rpm for 10 min. Subsequently, RBCs were fixed with 1% glutaraldehyde for 2 h and were washed thrice with PBS buffer, pH 7.4, by centrifugation at 3000 rpm for 10 min each. Following fixation and washing, a pool of RBCs was subjected to a dehydration process by sequential exposure to a series of ethanol concentrations (10%, 30%, 50%, 70%, 90%, and 100%) for 15 minutes each, followed by two sessions of dehydration, 30-minutes each, with 100% ethanol. A drop of RBCs was spread onto a glass coverslip, and it was allowed to dry overnight at room temperature. The dried RBCs were then coated with platinum and imaged using SEM (Carl Zeiss-SUPRA 55VP). The ability of NAC to neutralize the effect of pBQ on RBC morphology was assessed by incubating the RBCs treated with 0.5 mM pBQ alongside 0.5 mM and 1 mM NAC in separate sets.

### Estimation of free thiol groups of membrane proteins

The free thiol groups in the membrane proteins were quantified using DTNB kinetics as per the protocol of Y. Ando with minor changes [21]. Briefly, ghost RBCs suspension having 20 mg/mL membrane proteins were suspended in the PBS buffer, pH 7.4, and exposed to 1mM pBQ for 4 hr at 37 °C. The ghost RBCs were washed thrice with PBS, pH 7.4, buffer by centrifugation at 10,000 rpm for 10 minutes and then resuspended in PBS buffer, pH 7.4. The ghost RBCs were then treated with 100 µM DTNB for 15 minutes, followed by centrifugation at 10,000 rpm for 10 minutes. The supernatant comprising the TNB chromophore, a direct measure of thiol content, was then measured at 412 nm using a UV-Vis spectrophotometer (Shimadzu, UV-1900i).

### RBC membrane lipid peroxidation

Lipid peroxidation was determined by monitoring the Thiobarbituric Acid Reactive Substances (TBARS) levels, following the method described by Mandal *et al.* [15] with minor modifications. TBARS serves as a marker for the oxidative degradation of lipids, especially polyunsaturated fatty acids. TBARS forms through the interaction of thiobarbituric acid (TBA) with the byproducts of lipid peroxidation. RBCs (20% hematocrit) suspended in PBS buffer, pH 7.4, were exposed to 1 mM pBQ for 4 h at 37 °C, followed by washing and centrifugation at 3000 rpm for 10 min. Proteins were precipitated by the addition of 30% trichloroacetic acid (w/v), and the mixture was centrifuged at 6000 rpm for 10 minutes. The resultant supernatant was treated with 1% TBA (w/v) dissolved in 0.05 N NaOH and heated to 95 °C for 30 minutes. After cooling to room temperature, absorbance was measured at 532 nm using a UV-Vis spectrophotometer (Shimadzu, UV-1900i). The TBARS concentration was calculated using a molar extinction coefficient of 1.56 × 105 M^−1^ cm^−1^ [22].

### Measurement of RBC sedimentation

To assess the rate at which RBCs settle at the bottom, sedimentation of RBCs was measured in a container over a defined period, which provides insights into the rheological properties of blood. RBCs (10% hematocrit) were exposed to 1 mM pBQ for 4 h at 37 °C and were subsequently washed with PBS buffer, pH 7.4. Both pBQ-treated and untreated RBCs were then transferred into 1 mL syringes (DISPO VAN) with a final volume of 0.5 mL. The sedimentation rate was subsequently measured after a 6-hour period.

### NAC or GSH binding with pBQ

The interaction between GSH and NAC with pBQ, separately, was assessed by DTNB reaction and by determining the thermodynamic parameters using isothermal titration calorimetry (ITC) (in MicroCal PEAQ-ITC). For the DTNB reaction, 50 µM GSH or NAC was incubated with varying concentrations of pBQ for half an hour at 37 °C and subsequently exposed to 100 µM DTNB for 10 min. TNB chromophore was quantified at 412 nm using a UV-vis spectrophotometer (Shimadzu, UV-1900i). For ITC, 100 µM of NAC and GSH, separately, dissolved in the PBS buffer, pH 7.4, was loaded into the sample cell, and titration was carried out with 1 mM pBQ dissolved in the PBS buffer, pH 7.4. The experiment was performed by 2 µL consecutive injection with a 150 s interval. The heat change for each injection was plotted against the molar ratio of GSH and NAC, separately, with pBQ using Malvern MicroCal PEAQ-ITC control software of the computer attached to the instrument and was fitted to obtain the associated thermodynamic binding parameters.

## Results and discussion

### GSH estimation

GSH serves as an antioxidant that protects cells against oxidative damage. In the human body, the concentration of GSH across different tissues ranges from 1-10 mM. GSH reduces ROS, such as free radicals, through its oxidation to GSSG. GSH can be replenished using glutathione reductase in the presence of NADPH, thereby maintaining the GSH/GSSG ratio necessary for the cell to function [23]. Exposure to cigarette smoke has been shown to reduce GSH levels within RBCs [24,25]. In this study, we investigated the impact of pBQ on GSH levels in RBCs by subjecting them to varying concentrations of pBQ for 4 hours, as described in the methodology section. We observed a significant decline in the GSH levels from 1.13±0.04 mM in the control group, RBCs without the exposure of pBQ, to 0.098±0.01 mM following exposure to 0.25 mM of pBQ (Fig. 1A). GSH levels for 0.5 mM, 0.75 mM, and 1 mM pBQ exposed RBCs were found to be 0.0961±0.008, 0.088±0.012, 0.0654±0.007, respectively. This substantial decline in GSH level may be attributed to the covalent binding of pBQ with GSH (Fig. 1B) and the complex formed between GSH and pBQ (Fig. 1C), as characterized by the ESI-MS spectra. While exploring the effect of NAC on the GSH concentration, RBCs were treated with 0.5 mM and 1 mM NAC, separately, for 4 h at 37 °C, where we observed a significant increase in the concentration of GSH within RBCs across varying concentrations of NAC (Fig. 1D). However, upon incubating RBCs with varying concentrations of NAC and 0.5 mM pBQ, we observed that NAC was able to retain the GSH levels within RBCs (Fig. 1E). This suggests that NAC can neutralize pBQ. The possible mechanism of this neutralization is via covalent binding of NAC with pBQ, which was observed through DTNB reaction and ITC experiment (Supplementary Fig. 1). The thiol group of cysteine in NAC can act as a nucleophile, forming a covalent bond with the pBQ, thereby neutralizing the effect of pBQ.

**Fig. 1.**
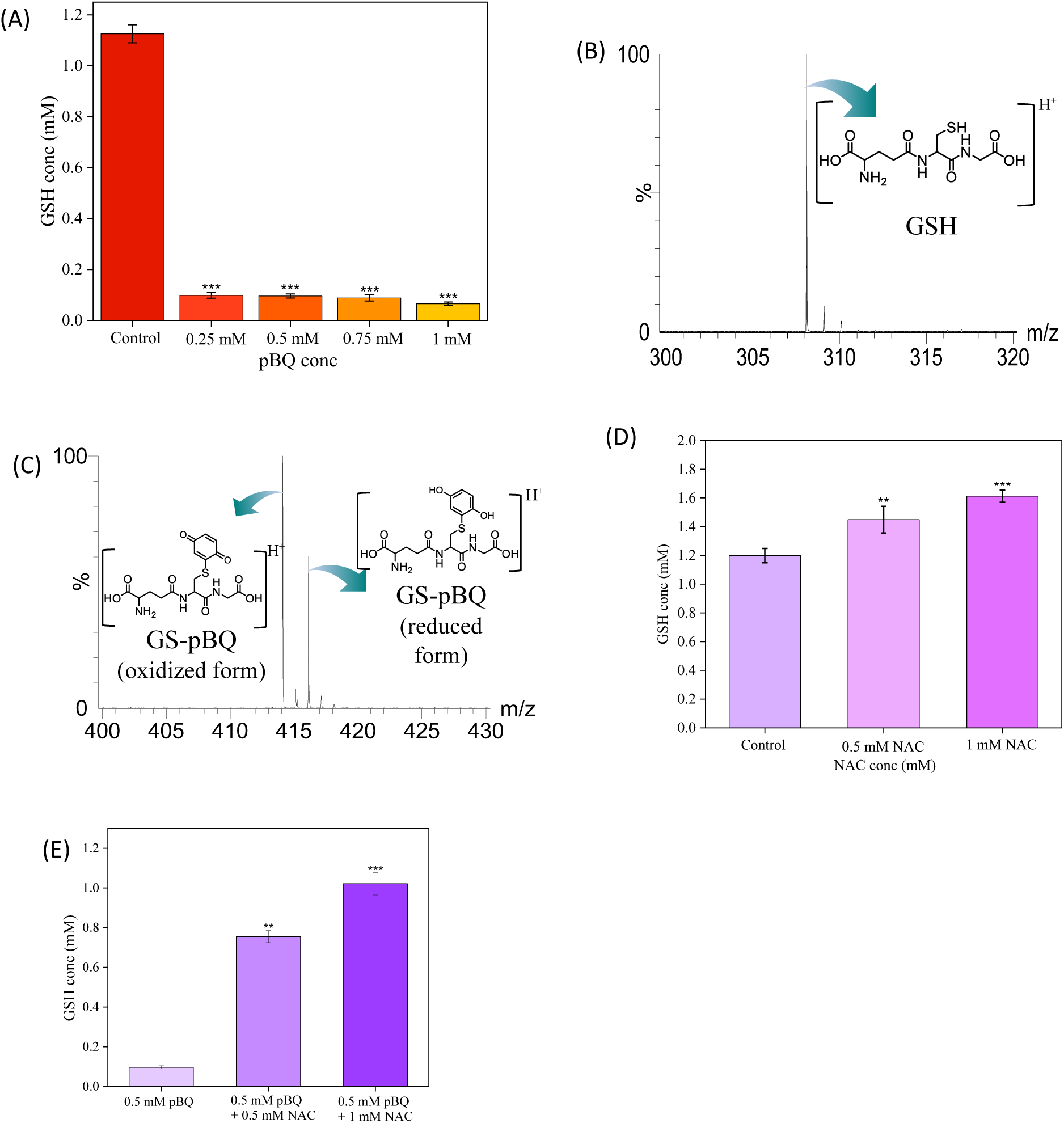
(A) GSH level of RBCs with and without (control) exposing them to varying concentrations of pBQ for 4 h, at 37 °C; (B) ESI-MS spectra of GSH; (C) ESI-MS spectra of the covalent complex between GSH and pBQ; (D) GSH levels of RBCs after exposing with varying concentration of NAC for 4 h at 37 °C; (E) GSH levels of RBCs after exposure to 0.5 mM pBQ alone and simultaneous exposure with 0.5 mM NAC, 1 mM NAC, in different sets for 4 h at 37 °C. The data are presented as the mean ± standard deviation of three replicates. *p ≤ 0.05, **p ≤ 0.01, ***p ≤ 0.001

The binding of GSH with pBQ is facilitated by the nucleophilic action of the thiol group of cysteine present in GSH (Scheme 1). The binding interaction of GSH with pBQ was demonstrated using the DTNB reaction and ITC experiment, as illustrated in Supplementary Fig.2. Moreover, the reduction in GSH levels might be exacerbated by its consumption in neutralizing the oxidants generated as a result of the pBQ exposure. The depletion of GSH levels might lead to an increase in ROS (Scheme 2), leading to oxidative stress, which can damage several cellular components, such as proteins, nucleic acids, and lipids, that might be associated with various pathological conditions such as cancer, COPD, etc [26–30].

**Scheme 1.**
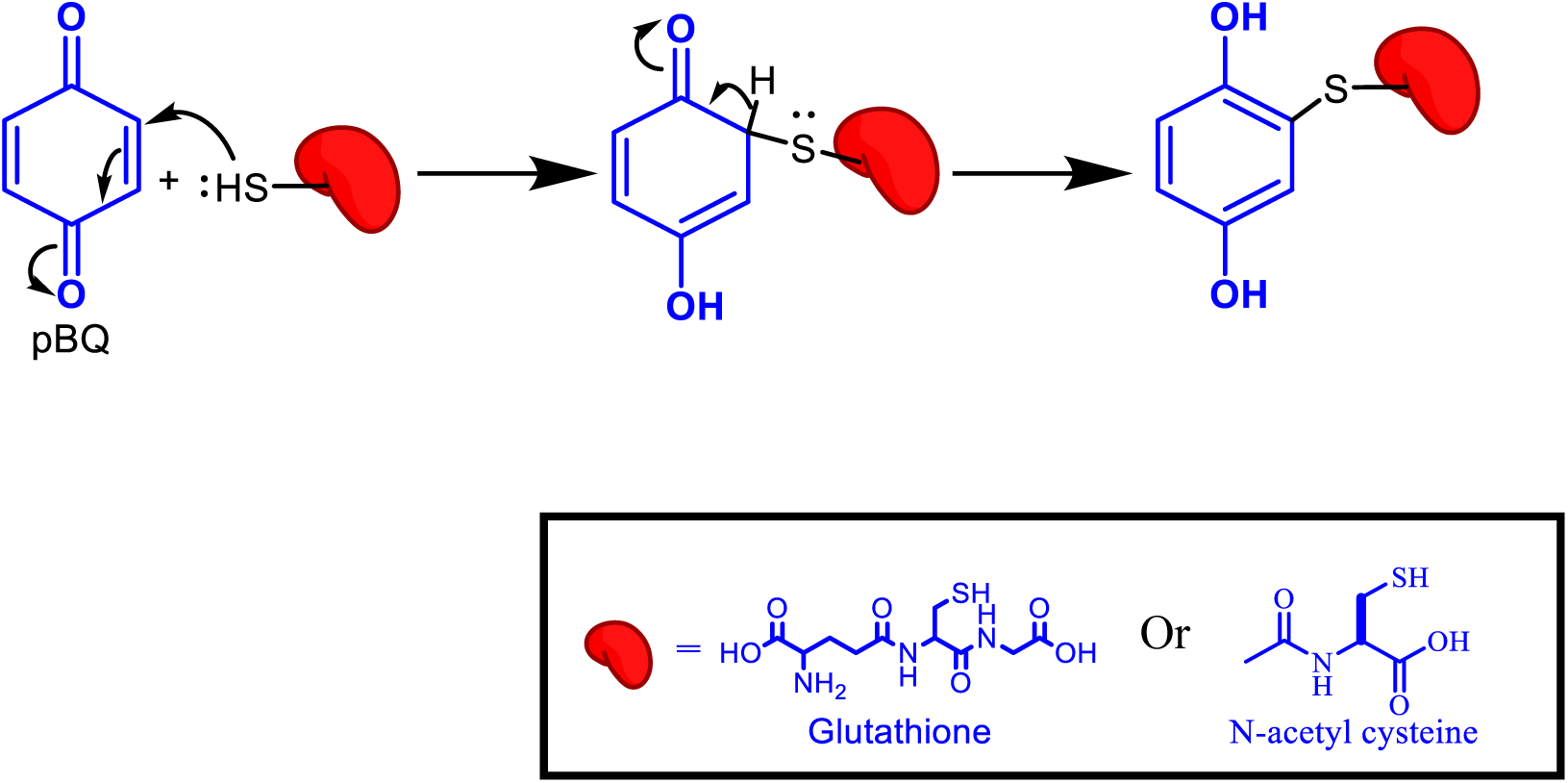
The mechanism of covalent binding of pBQ to the thiol (-SH) group of GSH/NAC.

**Scheme 2.**
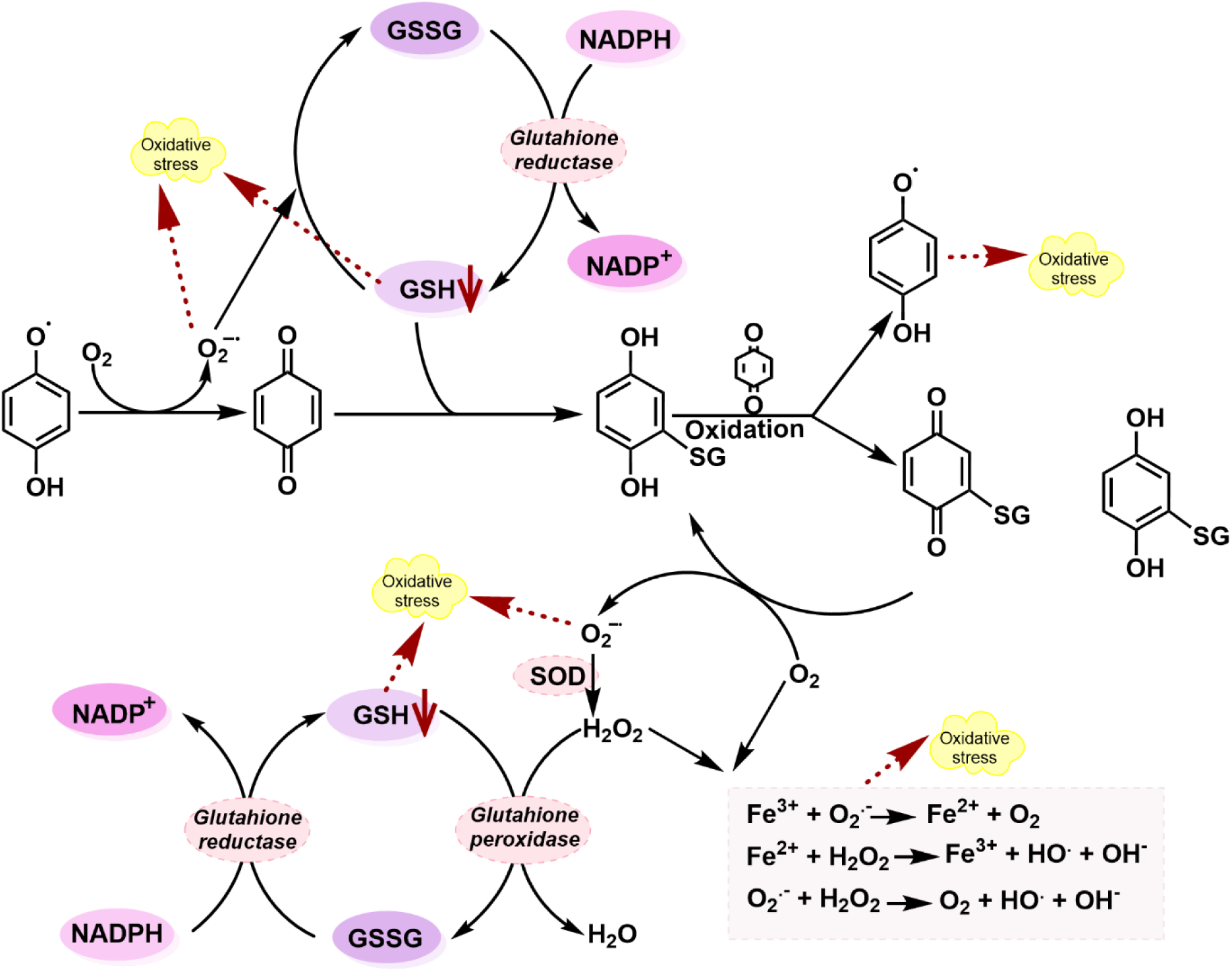
Different pathways lead to the generation of oxidative stress within RBCs following exposure to pBQ.

### ROS level measurement

ROS are the byproducts of cellular metabolism and play essential roles in various biological processes. However, excessive ROS production disrupts the balance between oxidants and antioxidants, leading to oxidative stress. ROS include both free radicals like hydroxyl radicals and non-radical intermediates like hydrogen peroxide, which can inflict oxidative damage on cells [31]. In our study, we investigated the impact of pBQ, a component of cigarette smoke, on RBCs by exposing them to different concentrations of pBQ. We observed a direct linear correlation between pBQ concentration and ROS levels, measured through fluorescence intensity originating from DCF (Fig. 2A). Furthermore, we found that GSH concentration decreased in the presence of pBQ, resulting in continuous generation of ROS without being neutralized. Cells maintain a definite level of antioxidants to counteract the generated ROS level; external oxidative stressors disrupt this equilibrium, leading to increased oxidative damage [32,33]. Cigarette smoke is known to increase ROS levels which has numerous downstream effects, including autophagy and altered mTOR levels [34,35].

**Fig. 2.**
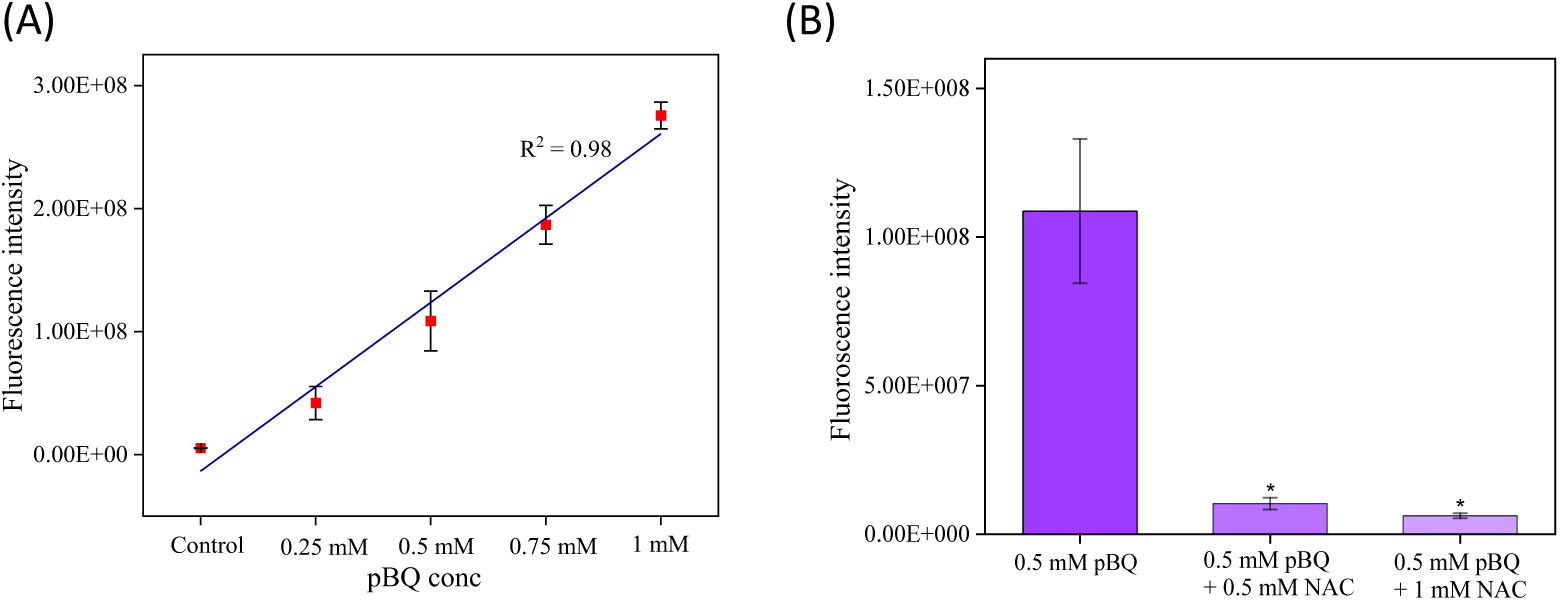
(A) ROS level of RBCs after 4 h of incubation with and without (control) varying concentrations of pBQ at 37 °C; (B) ROS level of RBCs with and without (only 0.5 mM pBQ) simultaneous incubation of 0.5 mM pBQ with 0.5mM NAC and 1mM NAC, separately, for 4 h at 37 °C; The data are presented as the mean ± standard deviation of three different experiments. *p ≤ 0.05, **p ≤ 0.01, ***p ≤ 0.001.

To assess the efficacy of NAC, an antioxidant, in mitigating pBQ-induced ROS generation, we incubated RBCs with NAC and pBQ concurrently. We observed that the ROS levels were approximately equal to the level of control (without pBQ exposure) in both 1:1 (pBQ: NAC) and 1:2 (pBQ: NAC) sets (Fig. 2B). Moreover, there was a significant reduction in ROS level within the RBCs exposed to the 1:1 and 1:2 of pBQ: NAC compared to RBCs solely exposed to 0.5 mM pBQ. This suggests that NAC effectively neutralizes pBQ, thereby reducing ROS generation.

### RBC membrane fluidity

The RBC membrane fluidity was assessed using the DPH, a fluorescent probe that preferentially embeds within the lipid bilayer of the cell membrane. The fluorescence of the probe is sensitive to the dynamic environment of the membrane lipids, making it a reliable indicator of membrane fluidity. Fluorescence anisotropy is inversely related to membrane fluidity and directly proportional to the microviscosity of the membrane. Fig. 3A shows the strong direct correlation between pBQ concentration and anisotropy. After exposure to different concentrations of pBQ, fluorescence anisotropy was found to be increased linearly, and membrane fluidity was found to be decreased linearly (Fig. 3A and 3B). The increase in microviscosity of the membrane due to alteration in the lipid bilayer might be responsible for decreased membrane fluidity. Previous studies have indicated decreased membrane fluidity in the RBCs of cigarette smokers, which is attributed to the changes in the phospholipid bilayer [19,20]. To elucidate the underlying mechanisms of altered membrane fluidity, we investigated the impact of pBQ on membrane lipids and proteins, fundamental constituents of the membrane.

**Fig. 3.**
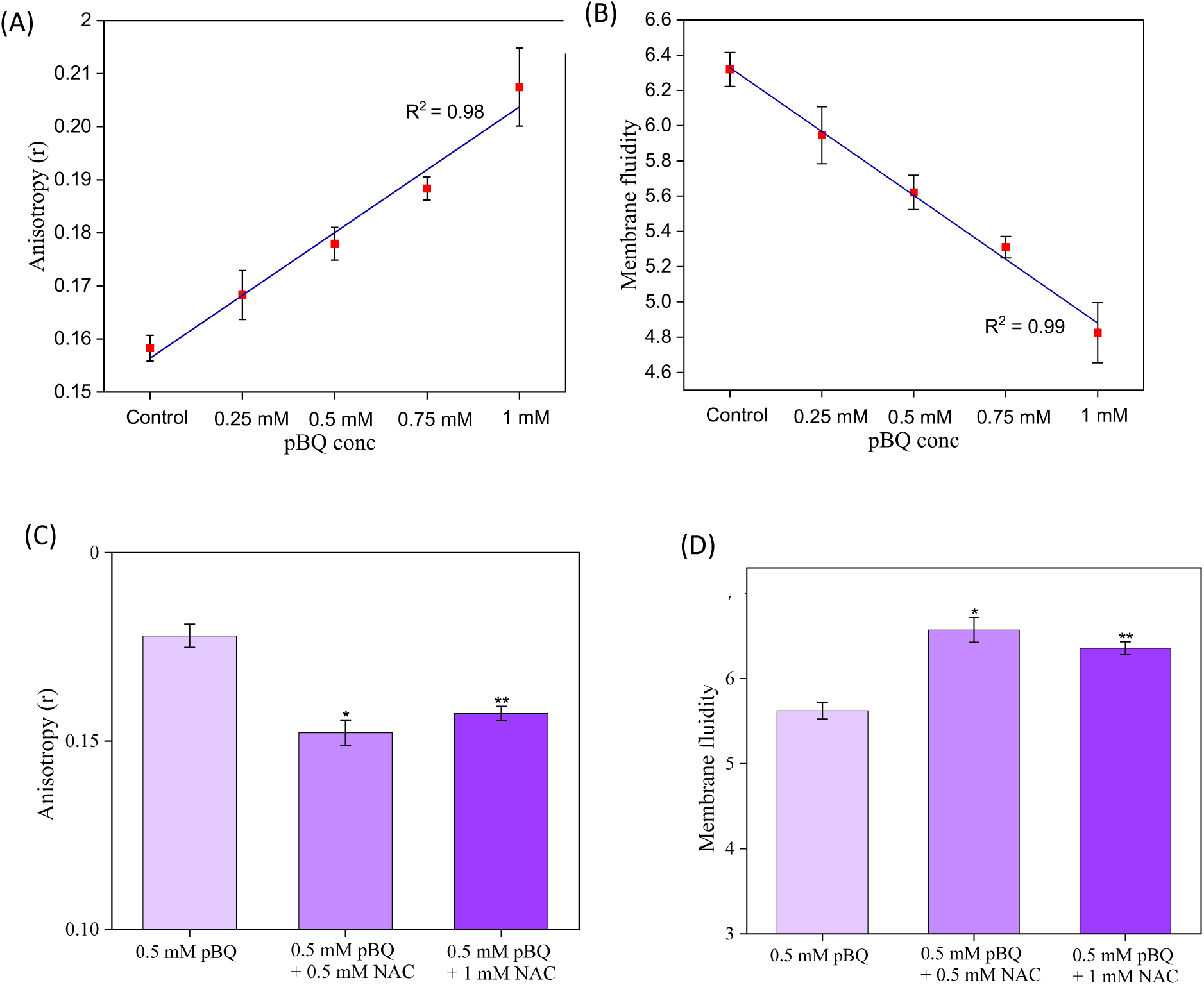
(A) Fluorescence anisotropy of ghost RBCs with and without (control) exposing them to varying concentrations of pBQ for 4 h, 37 °C; (B) Membrane fluidity of ghost RBCs with and without (control) exposing them to varying concentrations of pBQ for 4 h, 37 °C; (C) Anisotropy of ghost RBCs with and without (only 0.5 mM pBQ) simultaneous incubation of 0.5 mM pBQ with 0.5mM NAC and 1mM NAC, separately, for 4 h, 37 °C; (D) Membrane fluidity of ghost RBCs with and without (only 0.5 mM pBQ) simultaneous incubation of 0.5 mM pBQ with 0.5mM NAC and 1mM NAC, separately, for 4 h, 37 °C; The data are presented as the mean ± standard deviation of three different experiments. *p ≤ 0.05, **p ≤ 0.01, ***p ≤ 0.001.

The incubation of RBCs with pBQ in the presence of NAC indicates that NAC effectively neutralized the pBQ-induced alterations in RBC membranes. As shown in Fig. 3C and 3D, the concurrent incubation of RBCs with 0.5 mM pBQ and NAC at ratios of 1:1 and 1:2 resulted in unchanged fluorescence anisotropy and membrane fluidity, respectively, comparable to the control group of untreated RBCs. This indicates that NAC exerts a protective effect against pBQ-induced changes, likely by binding to pBQ and preventing its detrimental impact on membrane integrity.

### Free thiol estimation of RBC membrane proteins

Proteins are one of the important constituents of membranes and are involved in several biological functions. To assess the impact of pBQ on membrane proteins, we exposed ghost RBCs containing membrane proteins to 1mM pBQ for 4 h at 37 °C and observed a reduction in the concentration of the free thiol group of membrane proteins (Supplementary Fig. 3). The observed reduction in free thiol groups might be attributed to the direct interaction between pBQ and the free thiol groups present on RBC membrane proteins. Following pBQ exposure, this interaction likely contributes to the observed changes in membrane fluidity [36]. Additionally, membrane proteins are susceptible to oxidative damage with increased ROS production. Oxidation of proteins can result in several modifications, such as hydroxylation of aliphatic amino acid side chains and aromatic groups, sulfoxidation of methionine residues, nitration of aromatic amino acid residues, chlorination of aromatic groups, nitrosylation of sulfhydryl groups, and the formation of carbonyl adducts. These oxidative modifications can further lead to polypeptide chain fragmentation and the formation of protein aggregates through cross-linking [37]. Such alterations might also contribute to the disrupted membrane fluidity observed in the presence of pBQ-induced ROS generation.

### RBC membrane lipid peroxidation

Membrane lipid peroxidation was measured by quantifying TBARS, which represents the end products of lipid peroxidation that react with TBA. Exposure of RBCs to 1 mM pBQ resulted in a notable increase in TBARS production (Supplementary Fig. 4), indicating enhanced lipid peroxidation. This observation suggests that pBQ-induced elevation in ROS levels contributes to increased lipid oxidation. Polyunsaturated phospholipids are particularly susceptible targets for lipid peroxidation, which generates lipid hydroperoxides that degrade into various compounds utilized as markers of oxidative stress. This oxidative cascade can disrupt bilayer structure, altering membrane properties such as fluidity and compromising physiological functions, resulting in membrane damage [37]. Cigarette smokers exhibit a pronounced increase in lipid peroxidation, reflecting alterations in membrane organization or composition, with potential implications for various pathophysiological conditions [38,39]. Notably, smokers exhibit elevated levels of phosphatidylethanolamine and the corresponding decrease in phosphatidylcholine, factors that significantly impact membrane morphology and fluidity [40].

### RBC surface morphology

To assess the effect of the aforementioned alteration in the RBC membrane on its morphology, we have analysed RBCs through SEM. Exposure to varying concentrations of pBQ resulted in noticeable bulging and distortion on the surface of RBCs (Fig.4). These morphological distortions may be attributed to the peroxidation of membrane lipids, which disrupts lipid packing [37]. Similar morphological changes have been observed in the RBCs of smokers and patients with COPD [20,41,42]. To explore the potential neutralizing effect of NAC, we co-incubated RBCs with 0.5 mM pBQ and either 0.5 mM NAC (1:1) or 1 mM NAC (1:2) (Fig. 5) separately. The morphology of pBQ-exposed RBCs in the presence of NAC closely resembled that of untreated control RBCs, indicating a protective effect of NAC. This protection is likely due to the direct binding of pBQ with NAC, which neutralizes the harmful effects of pBQ.

**Fig. 4.**
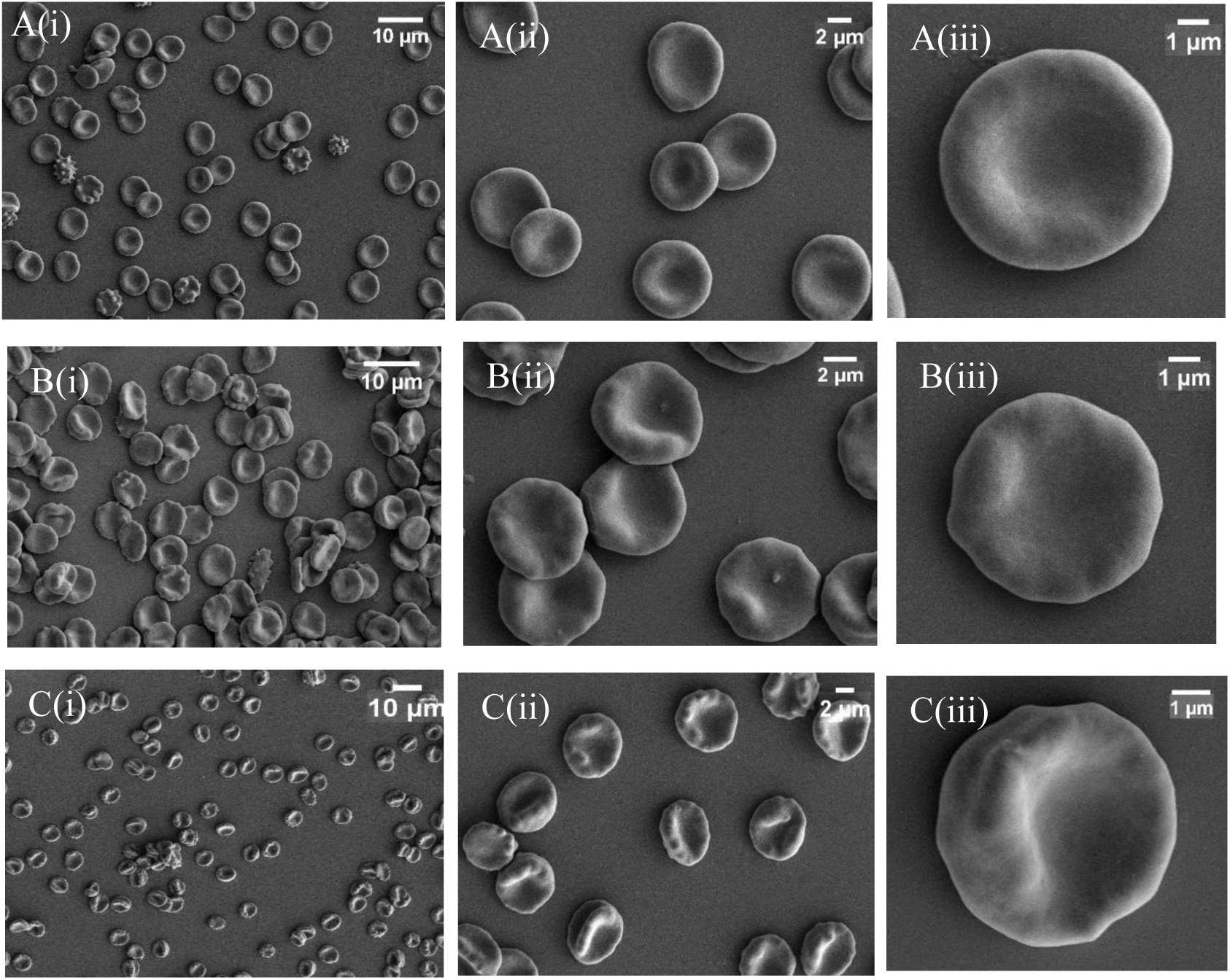
SEM-based analysis of the morphology of RBCs without pBQ (control) (A (i, ii, iii)) exposure for 4 h, 37 °C; (B (i, ii, iii)) with 0.5 mM pBQ, (C (i, ii, iii)) 1mM pBQ exposure at different magnifications.

**Fig. 5.**
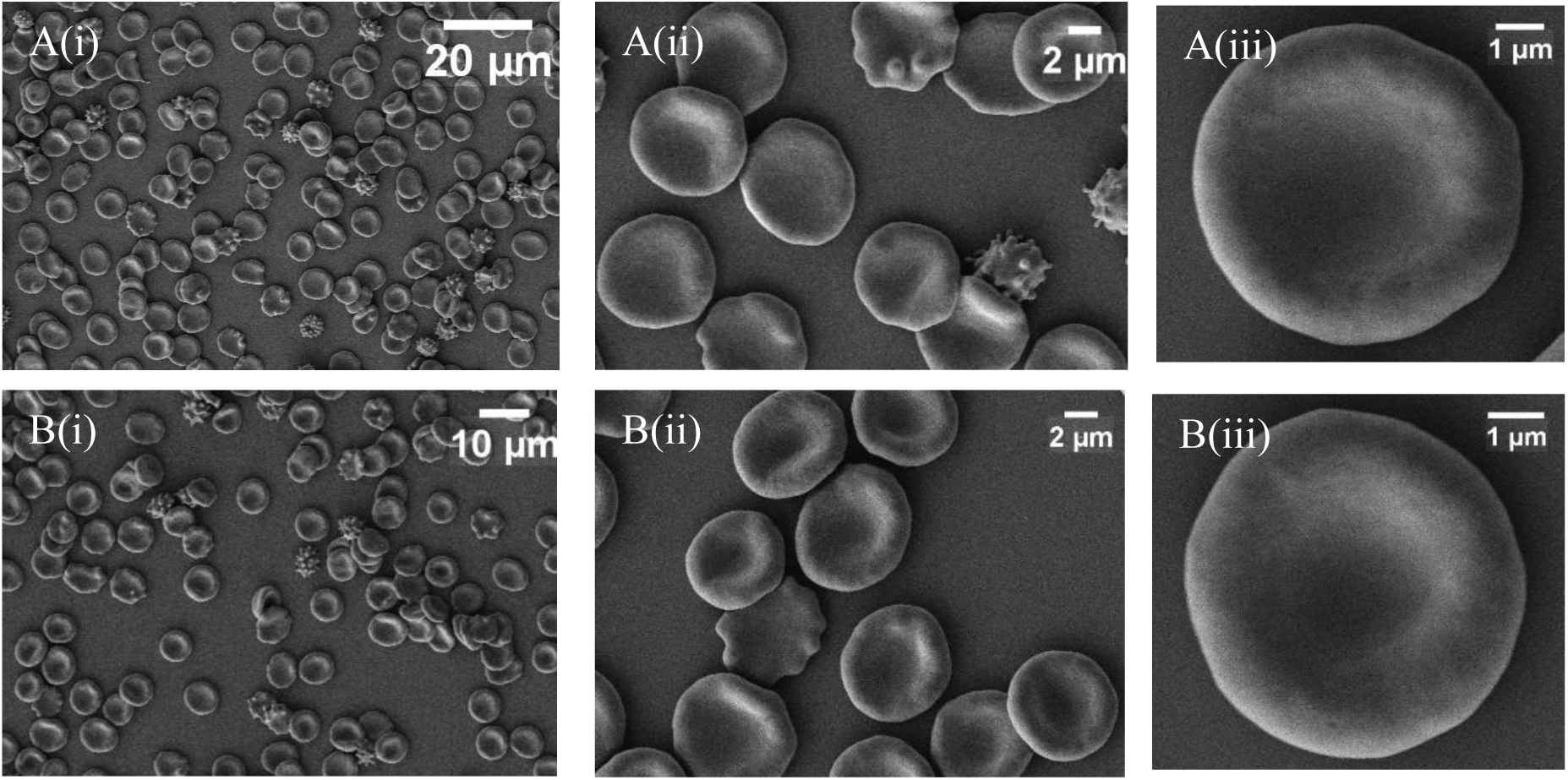
SEM-based analysis of the morphology of RBCs, after simultaneous incubation of 0.5 mM pBQ with 0.5 mM NAC (A (i, ii, iii)) and 1 mM NAC (B (i, ii, iii)), in separate sets for 4 h 37 °C, with different magnifications.

### Measurement of RBC sedimentation

The sedimentation of RBCs was assessed by measuring the volume of the packed RBCs, treated with or without pBQ for 4 h, after sedimentation for up to 6 h (Fig. 6A and 6B). The results revealed a lower packed cell volume in pBQ-treated RBCs compared to controls, indicating an increase in sedimentation in the case of pBQ treated RBCs. These findings suggest that the enhanced sedimentation reflects increased RBC aggregation, which might be due to the formation of larger clumps (rouleaux). Such aggregation might contribute to elevated blood viscosity and resistance to blood flow, potentially linking these changes in blood rheology to the cardiovascular disorders often associated with smoking.

**Fig. 6.**
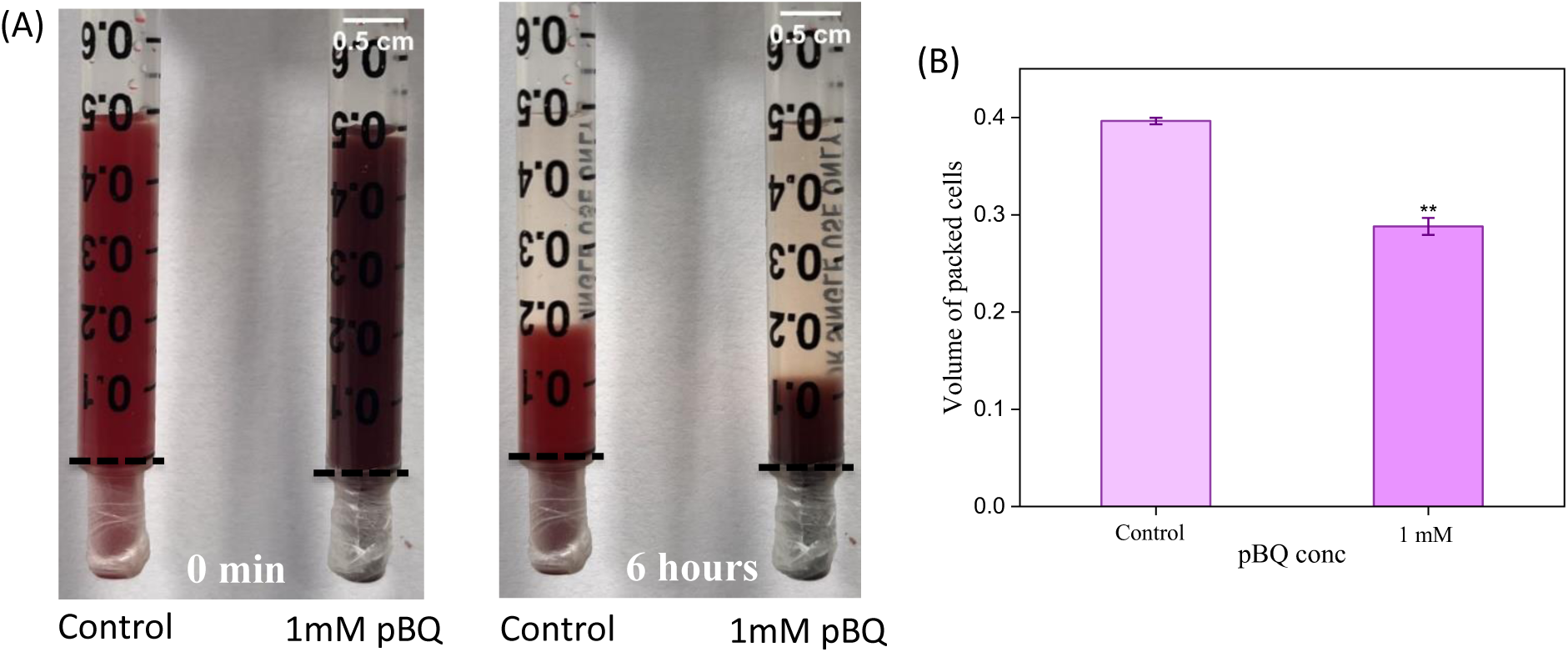
(A) RBC sedimentation after exposing them with and without (control) 1mM pBQ for 4 h, 37 °C followed by the sedimentation measurement for 6 hours in a 1 ml syringe.; (B) Volume of the packed RBCs after 6 h of sedimentation of RBCs exposed with and without (control) 1mM pBQ for 4 h, 37 °C. The data are presented as the mean ± standard deviation of three different experiments. *p ≤ 0.05, **p ≤ 0.01, ***p ≤ 0.001.

### GSH and NAC binding with pBQ

The binding interactions between NAC and GSH with pBQ were evaluated using two methods: DTNB kinetics and the ITC technique. In the DTNB kinetics assay, NAC/GSH was incubated with different concentrations of pBQ, revealing a linear decrease in TNB product formation, indicative of NAC/GSH consumption upon binding with pBQ (Supplementary Fig. 1A and 2A). Furthermore, the thermodynamic parameters for these interactions were determined using ITC.

ITC provides insights into binding interactions by elucidating thermodynamic parameters, including dissociation equilibrium constant (K_d_), changes in Gibbs free energy (ΔG), stoichiometry (N), and enthalpy change (ΔH). Supplementary Fig. 1B and 2B illustrate the ITC profile obtained from the titration of NAC and GSH, respectively, with pBQ at 37 °C. Each spike in the ITC profile corresponds to the heat change accompanying a single injection, with the intensity of spikes becoming uniform upon saturation of binding. Supplementary Fig. 1C and 2C depict the plot of heat change against the molar ratio of NAC and GSH, respectively, to pBQ, with the solid line representing the best-fit curve to the binding model. The data were fitted to a single-set binding sites model. Negative peaks in the data signify exothermic binding, and the calculated binding parameters are summarized in Table 1.

**Table 1.** Thermodynamic parameters derived from the ITC study for the binding of GSH and NAC, separately, with pBQ.

| Molecules | $K_d$ | $\Delta H$<br>(kcal/mol) | $\Delta G$<br>(kcal/mol) | $-T\Delta S$<br>(kcal/mol) | N |
| --- | --- | --- | --- | --- | --- |
| NAC | $219\text{e-}9 \pm 31.6\text{e-}9$ | $-22.4 \pm 0.234$ | -9.45 | 12.9 | $0.981 \pm 3.6\text{e-}3$ |
| GSH | $381\text{e-}9 \pm 39.4\text{e-}9$ | $-23.9 \pm 0.202$ | -9.11 | 14.8 | $0.736 \pm 2.9\text{e-}3$ |

The strong binding affinity between NAC/GSH and pBQ, as evidenced by the K_d_ values (219e-9 ± 31.6e-9 and 381e-9 ± 39.4e-9, respectively), emphasizes their substantial binding affinity and effective interaction. The negative ΔH and ΔG values depict the exothermic and spontaneous nature of the binding reaction, while the positive ΔS value suggests an increase in the randomness around the binding complex, indicative of an entropy-driven process.

The direct binding of GSH with pBQ elucidates its role in exacerbating oxidative stress by interacting directly with pBQ. This suggests that pBQ induces oxidative stress through two mechanisms, one by initiating a redox cycling cascade that ultimately leads to increased oxidative stress and the other by directly binding to GSH, which depletes intracellular antioxidant levels and consequently elevates oxidative stress. These dual pathways position pBQ as a potent inducer of oxidative stress, acting as an oxidizing agent through both redox cycling and antioxidant depletion.

Conversely, the neutralizing mechanism of NAC operates through two distinct ways: first, by directly binding to pBQ, thereby neutralizing its impact, and second, by bolstering the intracellular GSH levels, further reinforcing the antioxidant defense of the cell. The incubation of RBCs with pBQ in the presence of NAC demonstrates a pronounced neutralization effect, highlighting the potential therapeutic utility of NAC in mitigating pBQ-induced oxidative stress. This dual-action mechanism of NAC suggests its promising role as a protective agent against oxidative stress induced by toxins of cigarette smoke like pBQ.

## Conclusion

In conclusion, our study elucidates the significant impact of pBQ on various biochemical and morphological aspects of RBCs, highlighting its potential role as an oxidative stress inducer. Exposure to pBQ led to a marked decrease in GSH levels, increased ROS generation, and oxidative damage. Alterations in the RBC membrane, including changes in fluidity and morphology, further emphasize the detrimental effects of pBQ. These findings underscore the multifaceted nature of pBQ-induced oxidative stress and its potential implications for various pathological conditions. Although NAC can neutralize the effect of pBQ, the preincubated RBCs with NAC followed by pBQ exposure indicated that the modification of the cellular components executed by pBQ appears to be irreversible (Supplementary Fig. 5). By elucidating the molecular mechanisms underlying oxidative stress responses, our study contributes to advancing the understanding of cellular physiology in the context of environmental toxins like pBQ.

## Acknowledgment

Neha Yadav acknowledges the senior research fellowship provided by the CSIR (Council of Scientific and Industrial Research), Government of India. Additionally, we would like to acknowledge all the participants who provided samples for this study. We acknowledge the Nano Mission, Department of Science and Technology, for funding a mass spectrometry facility sanctioned under the project SR/NM/NS-1068/2015. We thank the DST-FIST (Project no: (SR/FST/LS-II/2017/93) for the common instrument facility at IISER Kolkata. We would also like to thank Pradip Kumar Tarafdar (Associate Professor) at IISER Kolkata for the access to the fluorescence spectrophotometer.

