## Supplementary data for "Effect of p-benzoquinone, a cigarette smoke-derived component, on the architecture and function of human RBC"

Supplementary Fig. 1. (A) The measurement of NAC concentration upon interaction with varying concentration of pBQ, *in vitro*, via UV-vis spectrophotometer; (B) ITC profile of the NAC binding with pBQ denoting the heat change while titrating NAC with pBQ; (C) Normalized heat change of ITC data in respect to the molar ratio of pBQ, where solid line represents the best-fit curve. The data was fitted into the single-site binding model.

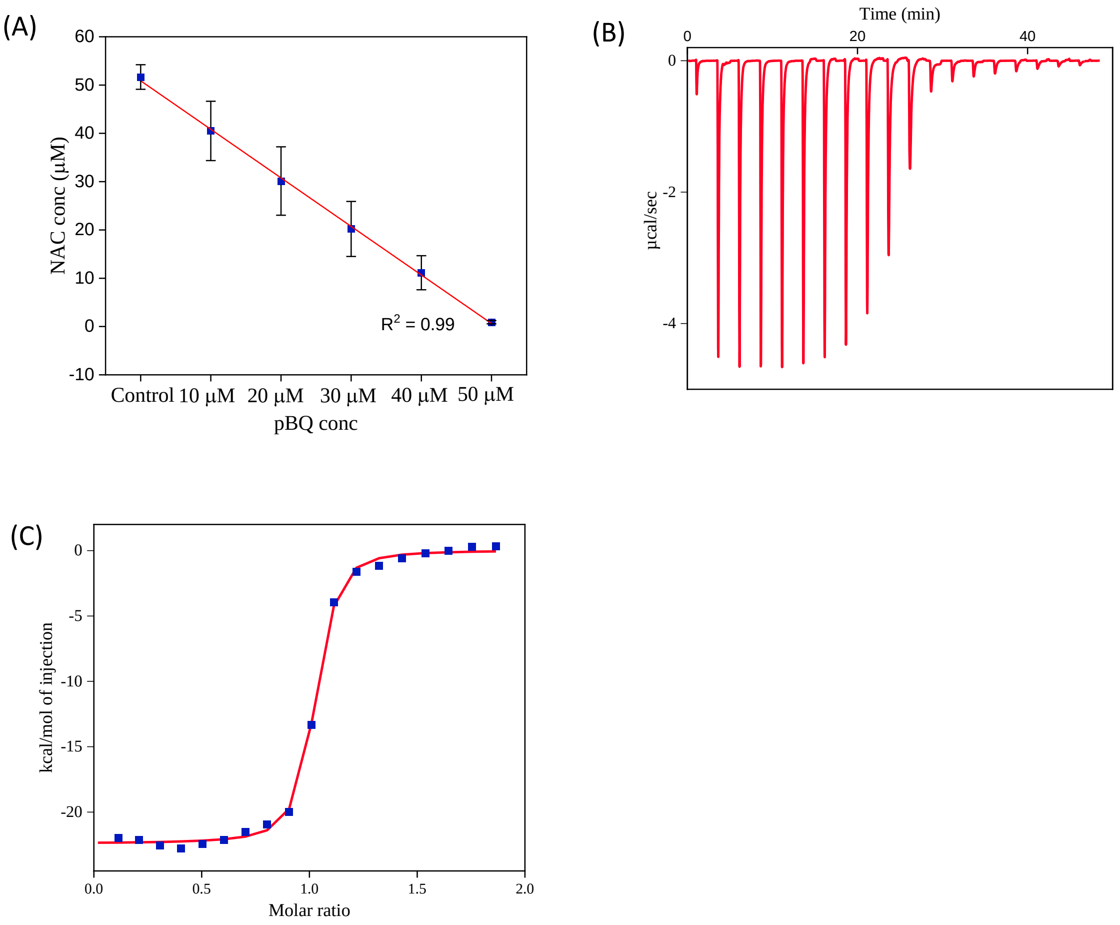

Supplementary Fig. 2. (A) The measurement of GSH concentration upon interaction with varying concentration of pBQ, *in vitro*, via UV-vis spectrophotometer; (B) ITC profile of the GSH binding with pBQ denoting the heat change while titrating GSH with pBQ; (C) normalized heat change of ITC data in respect to the molar ratio of pBQ, where solid line represents the best-fit curve. The data was fitted into the single-site binding model.

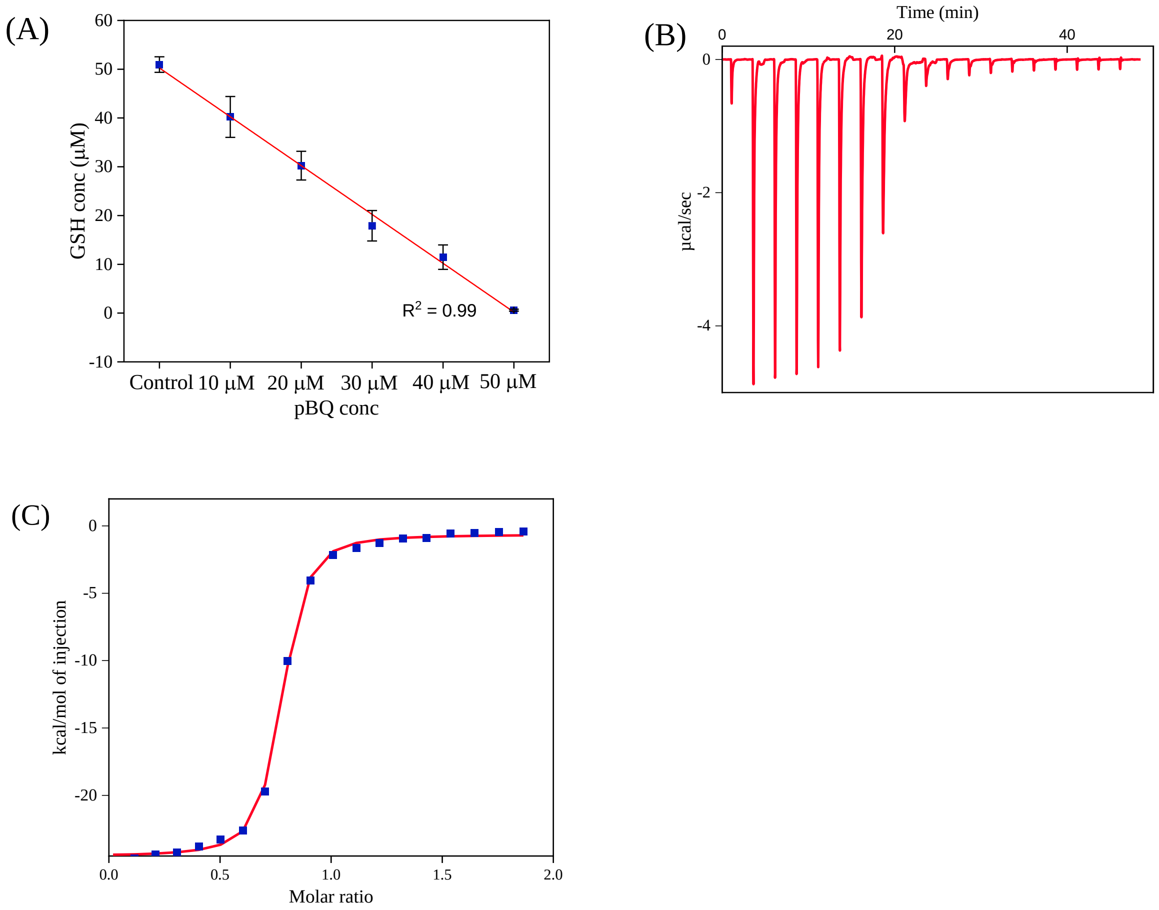

Supplementary Fig. 3. Quantified thiol group of ghost RBCs membrane proteins incubated with and without (control) 1 mM pBQ for 4 h, 37 ºC; The data are presented as the mean ± standard deviation of three different experiments. *$p \leq 0.05$, **$p \leq 0.01$, ***$p \leq 0.001$.

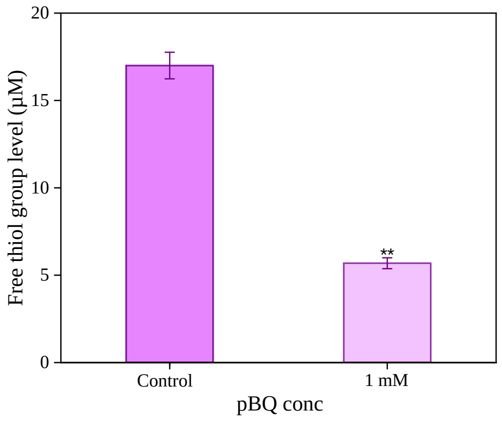

Supplementary Fig. 4. Quantified TBARS level of RBCs with and without (control) exposure to 1mM pBQ for 4 h at 37 ºC; The data are presented as the mean ± standard deviation of three different experiments. *$p \leq 0.05$, **$p \leq 0.01$, ***$p \leq 0.001$.

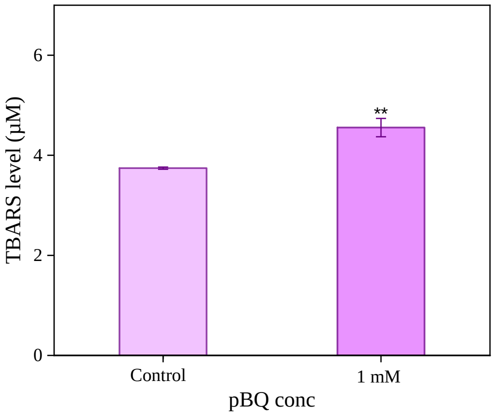

Supplementary Fig. 5. SEM-based morphological analysis of 0.5 mM pBQ exposed RBCs for 4 h at 37 ºC, followed by incubation with 0.5 mM NAC (A (i, ii, iii)) and 1 mM NAC (B (i, ii, iii)) in separate sets, for 4 h at 37 ºC, with different magnifications.

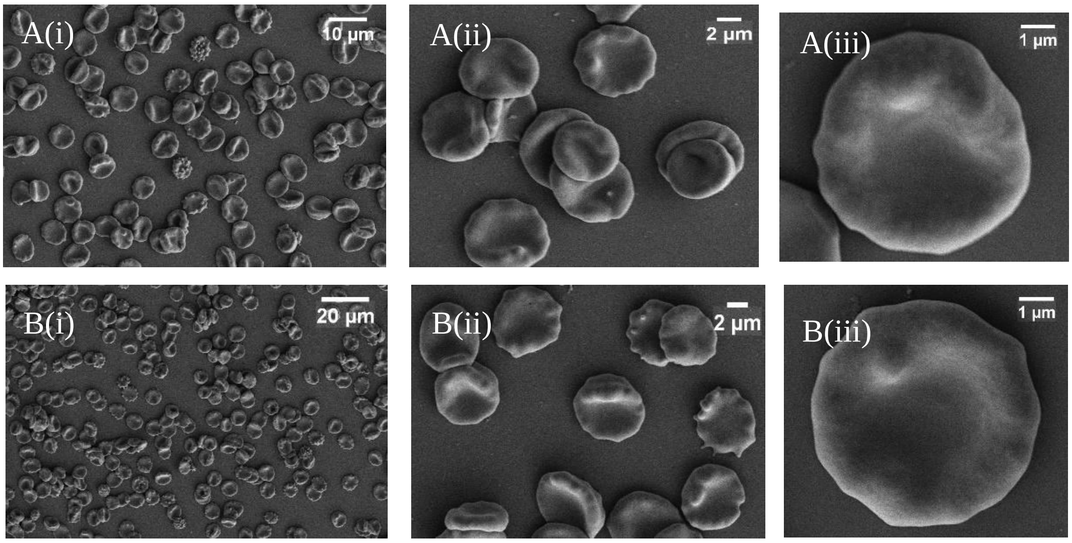
